# The Chromosome-based Genome of *Paspalum vaginatum* Provides New Insights into Salt-stress Adaptation

**DOI:** 10.1101/2022.08.08.503172

**Authors:** Li Liao, Xu Hu, Jiangshan Hao, Minqiang Tang, Longzhou Ren, Ling Pan, Shangqian Xie, Paul Raymer, Peng Qi, Zhenbang Chen, Zhiyong Wang, Jie Luo

## Abstract

Salinization is increasingly a major factor limiting production worldwide. Revealing the mechanism of salt tolerance could help to create salt-tolerant crops and improve their yields. We reported a chromosome-scale genome sequence of the halophyte turfgrass *Paspalum vaginatum*, and provided structural evidence that it shared a common ancestor with *Z. mays* and *S. bicolor*. A total of 107 *P. vaginatum* germplasms were divided into two groups (China and foreign group) based on the re-sequenced data, and the grouping findings were consistent with the geographical origin. Genome-wide association study (GWAS) of visually scored wilting degree and withering rates identified highly significant QTL on chromosome 6. Combination with RNA-seq, we identified a significantly up-regulated gene under salt stress, which encodes ‘High-affinity K^+^ Transporter 7’ (*PvHKT7*), as strong candidates underlying the QTL. Overexpression of this gene in *Arabidopsis thaliana* significantly enhanced salt tolerance by increasing K^+^ absorption. This study adds new insights into salt-stress adaptation of *P. vaginatum* and serve as a resource for salt-tolerant improvement of grain crops.

## Introduction

Salinization is increasingly a major factor limiting production worldwide. Saline alkali land accounts for 2.1% of the world’s total land and as much as 19.5% of irrigated land (www.fao.org/soils-portal). Most of the crops are salt-sensitive, which inhibits seedling growth and decreases yields when growing in salt-affected soils(Roy et al., 2014; Farooq et al., 2015; Acosta-Motos et al., 2017). Therefore, in the areas that have been already salt-affected or vulnerable to salinization because of seawater intrusion or storm surges, the development of salt-tolerant varieties is a very critical target.

Seashore paspalum (*Paspalum vaginatum*) has been utilized as turf for almost one hundred years(Wu et al., 2018; Qi et al., 2019) and the species is largely diploid (2n=2x=20). *P. vaginatum* is an important halophytic warm-seasoned perennial grass widely used on athletic fields, golf courses, and landscape areas in tropical and subtropical regions. It can be propagated rapidly by stolons and rhizomes to form a fine-textured turf(Liu et al., 2017; Wu et al., 2020). Owing to its tolerance to abiotic stresses, and its ecological aggressiveness, it is not only widely used as a turfgrass, but also serves as forage and ground cover for reducing erosion(Liu et al., 2017). *P. vaginatum* has strong salt tolerance and can even be watered with seawater for short periods making it especially important in locations near the sea or regions with water quality issues.

Because of the wide geographic distribution and long-term effects of natural or artificial selection, there are a number of diverse varieties for *P. vaginatum*, but only few genetic resources are currently available. For years, relevant studies have mainly focused on diversity analyses using random amplified polymorphic DNA (RAPD) markers(Liu et al., 1994), amplified fragment length polymorphisms (AFLPs)(Chen et al., 2005), and simple sequence repeat (SSR) markers(Shen et al., 2020), in attempts to identify loci associated with salt tolerance traits (Wu et al., 2018) or dollar spot resistance traits(Catching et al., 2019). A high-density genetic map of *P. vaginatum* indicated *P. vaginatum* is closely related to the important grain crops such as maize, sorghum and millets(Qi et al., 2019). Hence, greater understanding of the molecular mechanisms of *P. vaginatum* salt tolerance may provide a gateway opportunity to improve salt tolerance in cereal crops. Simultaneously it could be contributed as a model to the study of salt tolerance in grasses.

Here, we integrated Pacbio, Hi-C, RNAseq, Illumina short reads, and 10 × Genomics data, presenting a chromosome-scale assembly genome of *P. vaginatum* ‘SeaIsle2000’, a cultivar with high turf quality introduced to South China several years ago. To better understand the genetic basis of salt tolerance traits in *P. vaginatum*, we re-sequenced 107 accessions collected globally. By conducting a genome-wide association study (GWAS) and molecular experiment, we identified several QTLs and verified the *PvHKT7* gene related to salt resistance in *P. vaginatum*. Together, this study provided a foundation for accelerating the genetic improvement of *P. vaginatum*, furthermore this genetic resource may potentially be used for salt tolerance research and biotechnology assisted improvement of other grasses as well as other crops.

## Results

### Chromosome-scale Genome Sequencing and Assembly

A diploid cultivated *P. vaginatum*, SeaIsle2000, planted widely with high turf quality, was selected for genome sequencing, using four sequencing and assembly technologies: Illumina short-read sequencing, PacBio long-read sequencing, 10× Genomics and Hi-C data (Table S1). The genome size was estimated to be 589.20 Mb based on K-mer analysis (Fig. S1a) and flow cytometry analysis (Fig. S1b), close to the 600 Mb reported by previous estimates(Eudy et al., 2017). Primary genome contigs of *P. vaginatum* were produced by PacBio (∼94.90×). Then, we used Illumina reads (∼100.62×) to perform sequence error correction of contigs. The consensus sequences were further scaffolded by integrating with 10× genomics linked reads (∼92.52×). The assembly V1 consists of 185 scaffolds, and its contig N50 was up to 5.46 Mb while its scaffold N50 was 9.06 Mb (Table S2). We ploted the distribution of GC content and sequence depth by 10 kb slide window and found that this assembly had no GC bias and was polluted (Fig. S2). Finally, in order to improve the quality of assembly, 64.15 GB Hi-C clean reads were used to assist the assembly correction and mapped to raw genome assembly. After filtering those unmapped and duplicative reads, 117 scaffolds were clustered and ordered into 10 pseudo-chromosomes (Fig. S3), with a super-scaffold N50 of 47.06 Mb (Table S2, assembly V2).

The final *P. vaginatum* assembly captured 517.98 Mb of genome sequence, consisting of 10 chromosomes and 169 scaffolds (Table S2), with 485.64 Mb (∼93.76%) anchored into chromosomes (Table 1). We identified 1572 (97.40%) of the 1614 conserved genes with BUSCO and 239 (96.37%) of the 248 core eukaryotic genes with CEGMA in the genome assembly (Table S3). We found that 99.19% of the Illumina short reads mapped to the SeaIsle2000 genome. Furthermore, we used the 4078 SNP markers that were mapped on the first available published integrated genetic linkage map, the maternal parent map (HA map), the paternal parent map (AH map) and the both parents map (HH map), to validate scaffold assembly. 1854 markers (98.36%), 1637 markers (98.97%) and 1276 markers (98.91%) were hit on the HA map, AH map, and HH map, respectively (Table S4). Every linkage group largely corresponded to a single SeaIsle2000 chromosome, such as HA genetic map, but some chromosomes were completely inverted, such as chromosomes 2, 4, 5, 6 and 10 (Fig. S4). This showed that the chromosome sequences of the assembly are coincident with previous linkage groups. Taken together, these results suggest that the SeaIsle2000 genome is a high-quality assembly.

**Table 1.**
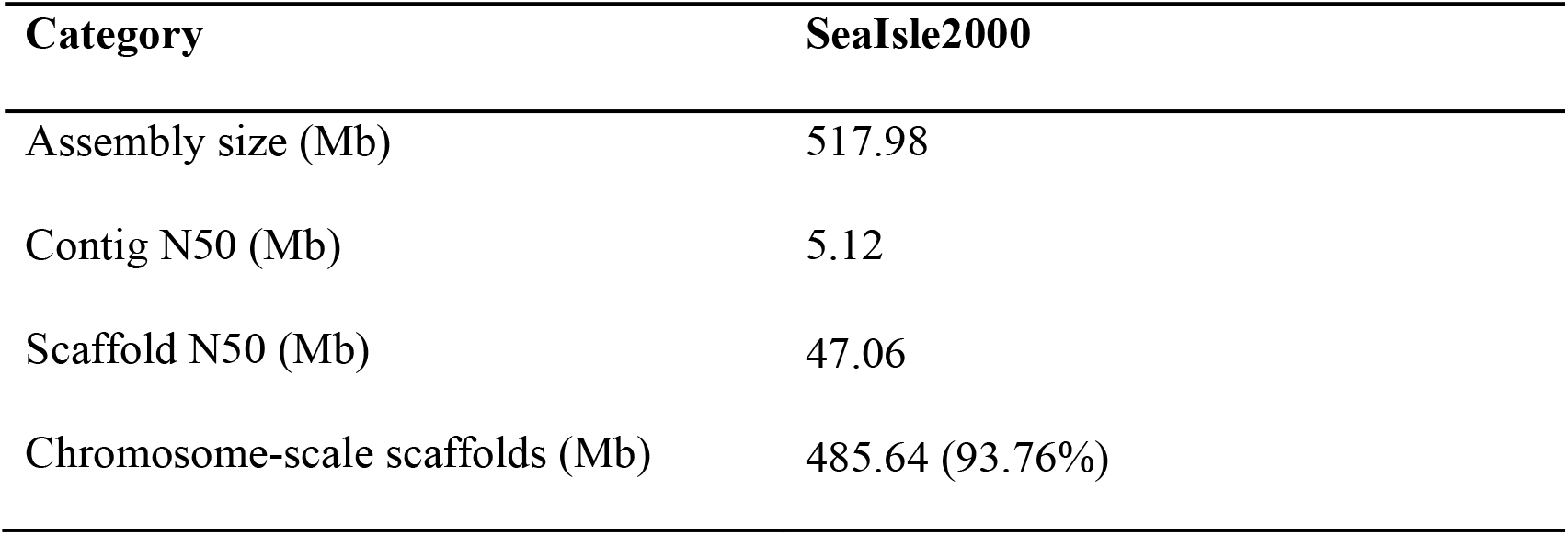
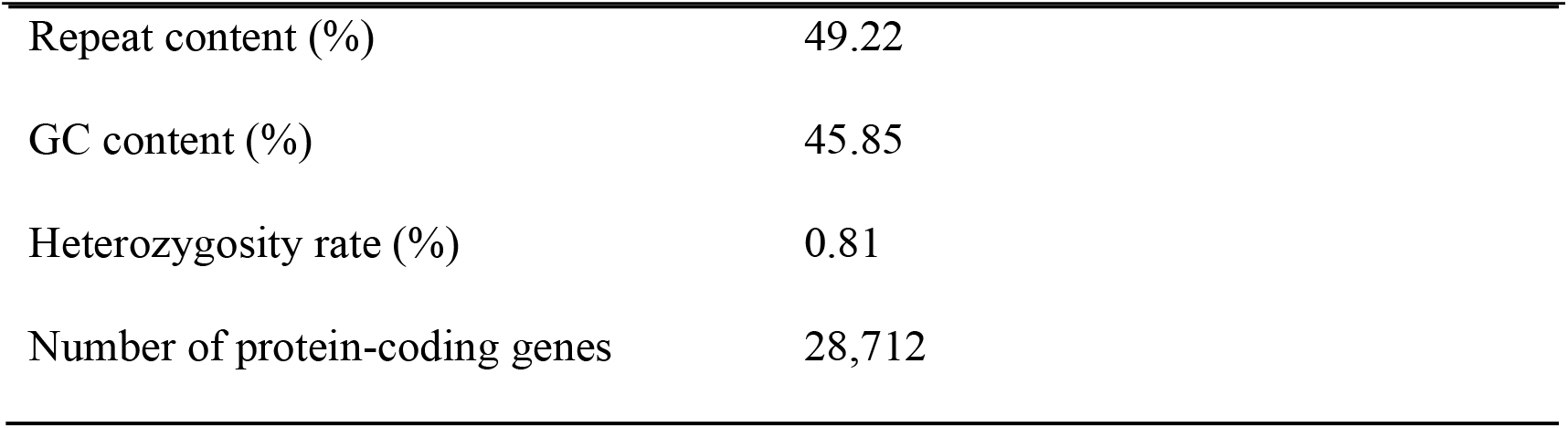
Summary statistics of the *P. vaginatum* genome assembly.

| Category | SeaIsle2000 |
| --- | --- |
| Assembly size (Mb) | 517.98 |
| Contig N50 (Mb) | 5.12 |
| Scaffold N50 (Mb) | 47.06 |
| Chromosome-scale scaffolds (Mb) | 485.64 (93.76%) |
| Repeat content (%) | 49.22 |
| GC content (%) | 45.85 |
| Heterozygosity rate (%) | 0.81 |
| Number of protein-coding genes | 28,712 |

### Genome Annotation

Based on de novo and homology-based predictions and transcriptome data, a total of 28,712 protein-coding genes were predicted, with an average coding sequence length of 3.53 kb and an average of 4.99 exons per gene, its gene structure was similarly to the common species, such as *A. thaliana, S. bicolor* and *Z. mays* (Table S5). Of the 28,712 predicted genes, 27,536 (95.90%) could be matched in at least one public protein database and given function descriptions (Table S6). As expected, gene density was higher on the arms than in the middle of the chromosomes (centromere regions) (Fig. 1a).

**Fig. 1.**
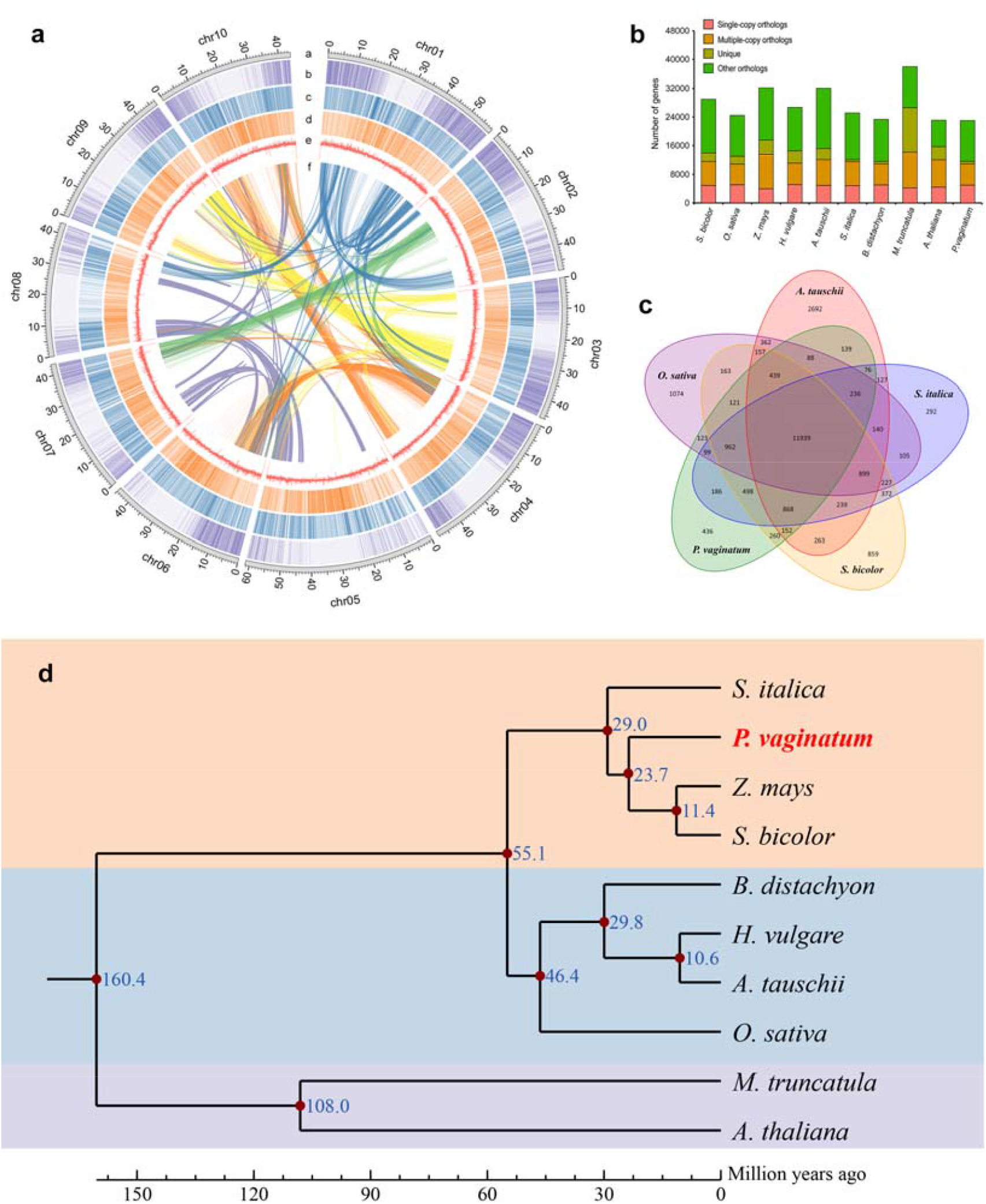
Genome evolutionary history. **a** Chromosomal features and synteny landscape of *P. vaginatum* genome. a, Chromosome size with units in Mb. **b**, Gene density. c, Indel density. d, SNP density. e, GC content. f, Genome syntenic blocks are illustrated with colored lines. b The distribution of single-copy, multiple-copy, unique, and other orthologs in the 9 plant species. **c** Venn diagram represents the shared and unique gene families among five species. Each number represents the number of gene families, numbers in the non-overlapped circlerepresent the number of gene families were specific to the species. **d** Phylogenetic tree of 10 plant species. Blue numbers represent divergence time of each node. The divergence time between the monocots and eudicots were estimated to be 138.2-190.0 million years ago (Ma).

There were 2,725 non-coding RNAs (555 miRNAs, 603 tRNAs, 331 rRNAs and 1236 snRNAs) predicted in the SeaIsle2000 genome sequence with a total length of 362.71 kb (Table S7). Curated repeat libraries and *de novo* prediction were used to annotate the repetitive elements. A total of 474,904 repetitive elements were identified, of which, 114,750 were tandem repeat elements with a mean length of 336.71 bp (Table S8).

### Comparative Genomic and Evolutionary Analysis

We applied the predicted proteomes of *P. vaginatum* and 9 other sequenced species to identify putative orthologous gene clusters. A total of 31,619 orthologous gene families composed of 338,128 genes were identified from 10 plant species of which, 6,949 clusters of genes were shared by the 10 species. 1800 (25.9%) of the shared gene clusters were single-copy families (Fig. 1b). Further, 11,939 gene families were present across *P. vaginatum, S. bicolor, O. sativa, S. italica* and *A. tauschii*. Compared with these four plant species, *P. vaginatum* genome contained 436 specific gene families containing 849 genes (Fig. 1c; Table S9). Gene Ontology (GO) term enrichment analyses of *P. vaginatum* specific genes showed that the cellular component termed ‘cytoplasmic stress granule’ was enriched; ‘cellular response to light intensity’, ‘cellular response to UV’ and ‘root hair elongation’ were enriched in the biological process categories (Fig. 2a; Table S10) which may be adapted to tropical climates in *P. vaginatum*. KEGG pathway enrichment analyses of *P. vaginatum* specific genes showed that ‘Linoleic acid metabolism’, ‘Flavonoid biosynthesis’ and ‘MAPK signaling pathway’, etc. were significantly enriched (Fig. 2b; Table S10) which may contribute to the special characteristic of salt-resistant in *P. vaginatum*.

**Fig. 2.**
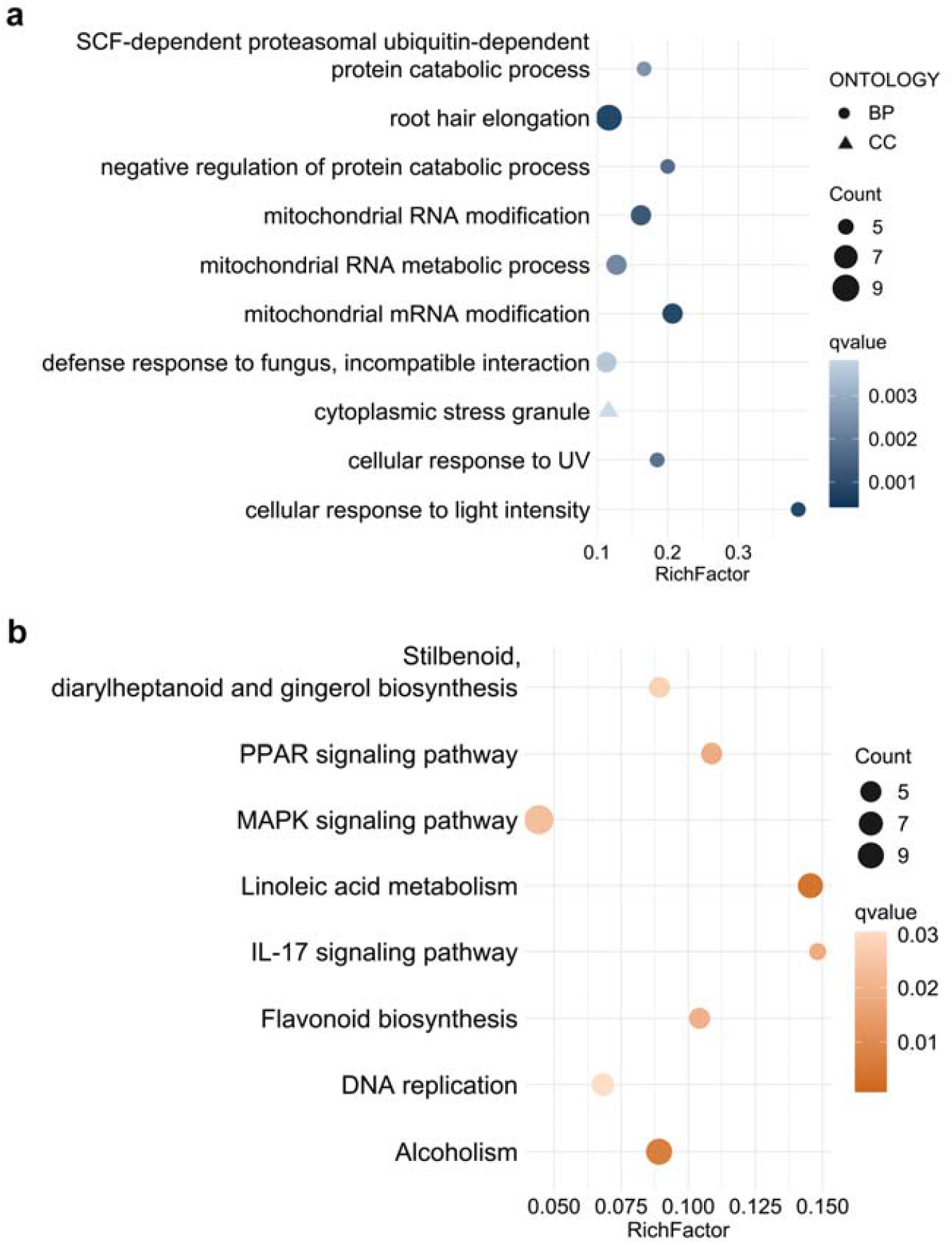
Enrichment analysis of specific genes in *P. vaginatum*. **a** The enriched GO terms with top 15 RichFactor were presented. The color of circles represents the statistical significance of enriched GO terms. The size of the circles represents the number of genes in a GO term. **b** The enriched KEGG pathways with corrected qvalue < 0.05 were presented. The color of circles represents the statistical significance of enriched KEGG pathways. The size of the circles represents the number of genes in a KEGG pathway.

A set of single-copy ortholog genes of the 10 plant species that included 8 monocots and 2 eudicots, and applied to construct a phylogenetic tree. Phylogenetic tree revealed *A. thaliana* and *M. truncatula* were clustered in one branch, *B. distachyon, H. vulgare, A. tauschii* and *O. sativa* were clustered in one branch, *S. italica, S. bicolor, Zea mays* and *P. vaginatum* were clustered in the other branch (Fig. 1d). These topologies of these species were consistent with the results of published papers. The divergence time between the monocots and eudicots were estimated to be 138.2-190.0 million years ago (Ma) (95% confidence interval). The tree indicated that *P. vaginatum, S. bicolor* and *Z. mays* might share a common ancestor and their divergence time was estimated to be about 23.7 Ma (the interval was 21.1-26.8 Ma).

According to the close relationship of *P. vaginatum, S. bicolor* and *Z. mays*, we investigated the whole-genome duplication (WGD) events during the evolutionary course of the 3 species. A total of 2,123 gene pairs were identified as segmental duplications in ten chromosomes of the *P. vaginatum* genome (Table S11). Among these gene pairs, 318 (14.98%) were located on the same chromosome, the others were across different chromosomes. The widespread gene duplications suggested that a WGD might have occurred during *P. vaginatum* genome evolution. Based on the previous research, there is a very well syntenic gene between *P. vaginatum* and *S. bicolor*. Here, more than half (60.45%) of the predicted *P. vaginatum* genes were identified as syntenic genes with *S. bicolor* genes, resulting in total 22,345 pair genes (Fig. 3a; Table S12). The synteny analysis between the two genomes provided clear structural evidence of a common ancestor. Further, we investigated the collinearity between *P. vaginatum* and *Z. mays*, the results showed that 16,046 *P. vaginatum* genes had good collinearity with *Z. mays* genes. Interestingly, many *P. vaginatum* genes have one-versus-four syntenic genes to *Z. mays* (Fig. 3b). The above results indicated that *P. vaginatum, S. bicolor*, and *Z. mays* might share a common ancestor, and *Z. mays* may have experienced two WGD events when compared with *P. vaginatum* (Fig. 3c).

**Fig. 3.**
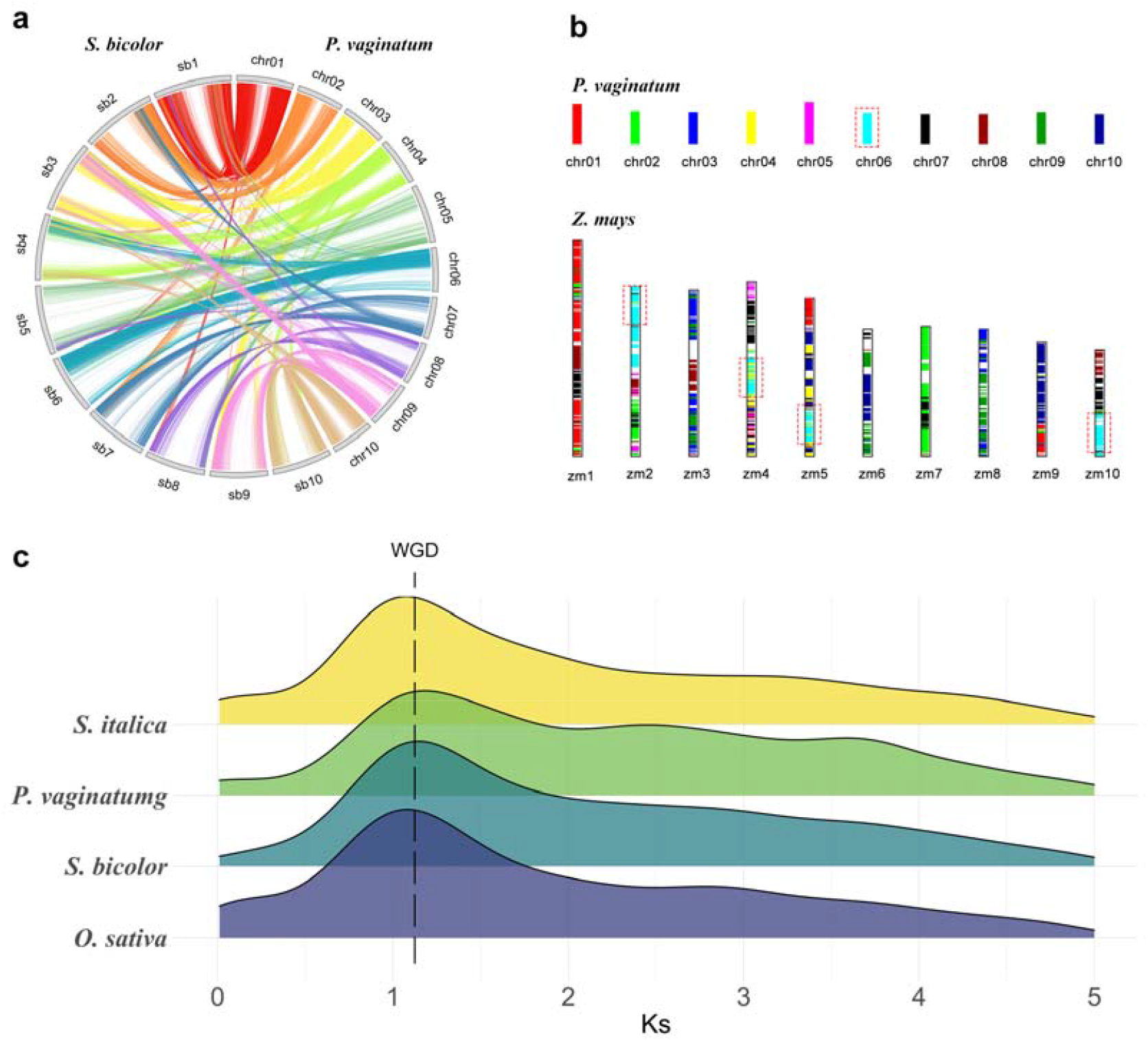
syntenic genes of *P. vaginatum, S. bicolor, Z. mays*. **a** the relationship of syntenic genes of *P. vaginatum* and *S. bicolor*. **b** the relationship of syntenic genes of *P. vaginatum* and *Z. mays*. Different colors mean different chromosomes in *P. vaginatum* genome, the corresponding color in *Z. mays* chromosomes mean that genes are syntenic with those genes located on corresponding chromosomes in *P. vaginatum* genome. Red dotted boxes show a case of the genes in chromosome 6 of *P. vaginatum* that have four copies in *Z. mays*. **c** Distribution of synonymous substitution rates (Ks) among collinear paralogs in four plants.

### Population structure and genomic variation

To explore genetic variation in *P. vaginatum*, we re-sequenced 107 germplasms with an average depth of 12.78× and 98.12% mapping rate of the SeaIsle2000 genome (Table S13). Using this dataset, we identified 3,083,910 high-quality SNPs and 1,023,374 InDels, corresponding to 6.35 SNPs and 2.11 InDels per kb (Table S14). A total of 68,402 SNPs (1.83%) and 19,196 InDels (1.45%) were located in gene coding regions (Table S15). Next, we investigated the genetic structure of the *P. vaginatum* population for clusters (K) from 2 to 10 based on 3.08 million SNPs. Delta K showed a peak at 2 (Fig. S5), suggesting two clusters as the most appropriate option. This supports the reliable result of two major discrete clusters of China and other countries from population the phylogenic tree and PCA (Fig. 4). But several germplasms were not well separated into two populations, indicating the occurrence of genetic diversity along with their adaptation to different environments. The China population of *P. vaginatum* had higher nucleotide diversity (π=2.04×10^−3^) than the Foreign population (π=1.06×10^−3^; Fig. S16). However, linkage disequilibrium (LD) decayed faster in the Foreign population than in China population (Fig. S6). Tajima’s D results of all germplasms was 1.86, while it was 2.67 and 1.55 in China and the other countries group, respectively, when all accessions were separated into different groups. This indicates that there are more alleles of high/medium frequencies in the population due to equilibrium selection and bottleneck effects. These results revealed that the allelic diversity in the China population was higher than that in the other countries population. FST between the two populations (FST = 0.37) demonstrated relatively higher genetic distance (Table S16).

**Fig. 4.**
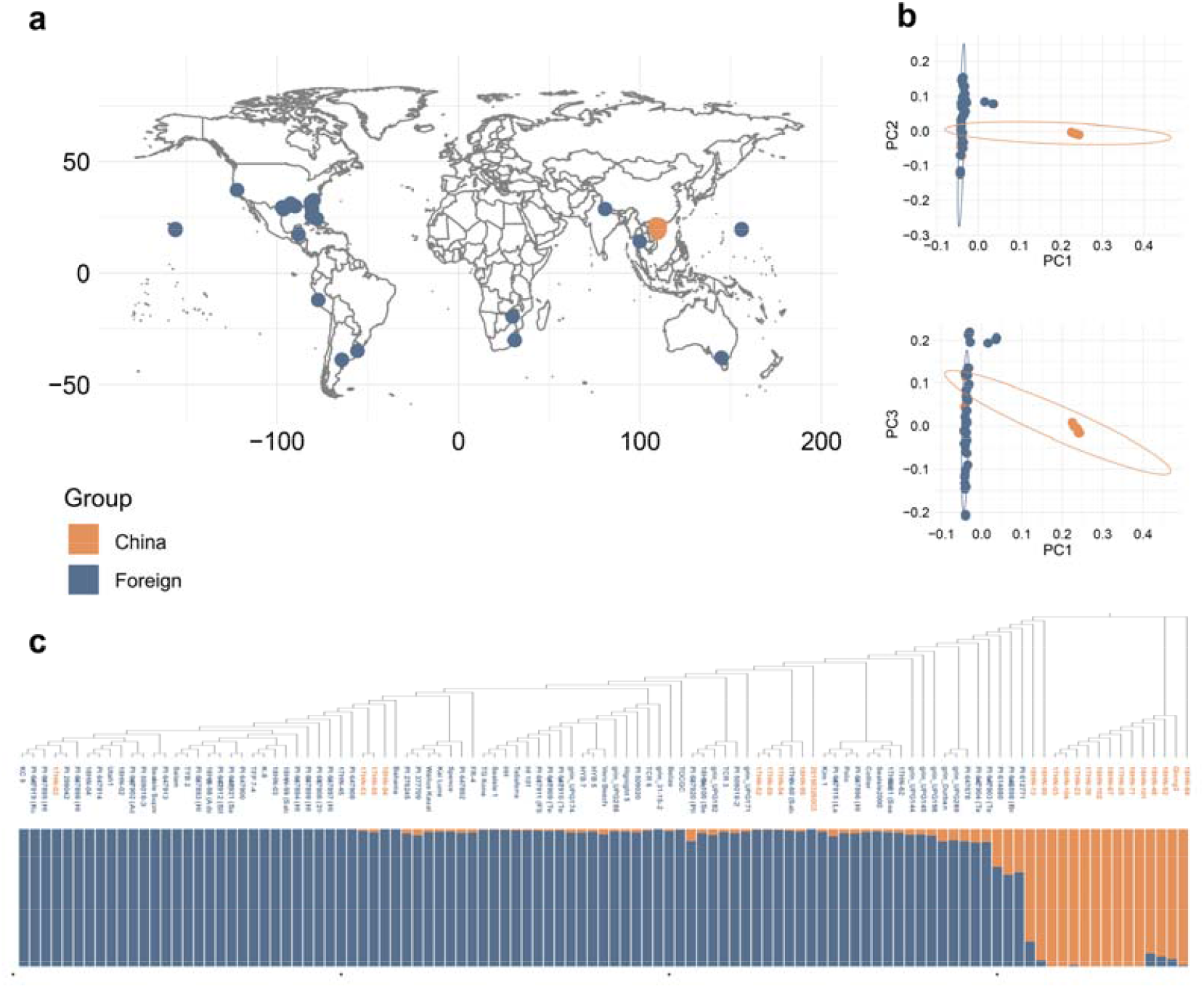
Geographical distribution and population structure of *P. vaginatum* accessions. **a** Geographic distributions of 107 *P. vaginatum* accessions. **b** PCA plots showing two divergent clades of 107 *P. vaginatum* accessions. **c** The NJ phylogeny of 107 *P. vaginatum* accessions and model-based clustering with K = 2.

### Population phenotypic variation under salt stress

The symptoms of plant leaves were observed and recorded under salt stress. We used two criteria, wilting degree (WD3, WD7) and withering rates (WR3, WR7), to estimate salt tolerance levels when evaluating the salt tolerance among 77 samples. The descriptive statistics of WC and WR are provided in Table S17 and the distribution for each trait is shown in Fig. S7a. The variable coefficients of WD3 and WD7 were 69.95% and 20.65%, respectively, and variable coefficients of WR3 and WR7 were 51.06% and 43.13%, respectively. These results indicated that a wide range of phenotypic variation among *P. vaginatum* germplasms were associated with salinity tolerance. The correlations between growth response and stress were obviously positive (0.33 to 0.80) and WR3 was highly correlated with WR7 (0.80). Furthermore, correlation analysis (Fig. S7b) showed that these 4 traits were significantly correlated with each other. We therefore adopted all of the traits as a meaningful indicator of salt tolerance.

### GWAS for salt tolerance

To identify candidate genes related to salt stress, annotations of the genes were analyzed within the identified loci (10.9 kb up and down stream of the most significant SNP), the *P*-value thresholds were set at 1.69×10^−7^ (significant, 0.5/n, -log_10_(P) = 6.77) and 1×10^−6^ (suggestive, -log_10_(P) = 6). Totally, 19 SNPs associated with resistance to salt were identified and 26 candidate genes were detected. Manhattan and QQ plot of GWAS results were shown in Fig. S9. These candidate genes distributed on chromosomes 3, 6, 9 and 10, with detailed information of the genes listed in Table S18. GO enrichment analysis was carried out to elucidate the specific biological functions of the 26 candidate genes. The significantly enriched GO terms concerning sodium ion transmembrane transport (GO:0035725), monovalent inorganic cation transport (GO:0015672), cation transmembrane transport (GO:0098655) etc (Table S19). Interestingly, a peak strongly associated with salt tolerance (Fig. 5a;5b), identified on chromosome 6, was located in emOS140.194 (*PvHKT7*), which was the orthologous gene with *HKT7*, a high-affinity potassium transporter in the *S. bicolor* genome. Five SNPs with the lowest P values on *PvHKT7* gene generated two haplotypes (Fig. 5c). And the five SNPs significantly associated with salt tolerance traits explained 46.98% ∼ 55.74% of the phenotypic variance (Table S20). The accessions with *PvHKT7*-Hap2 have better salt tolerance (*P*<0.05) than those with *PvHKT7*-Hap1 (Fig. 5d).

**Fig. 5.**
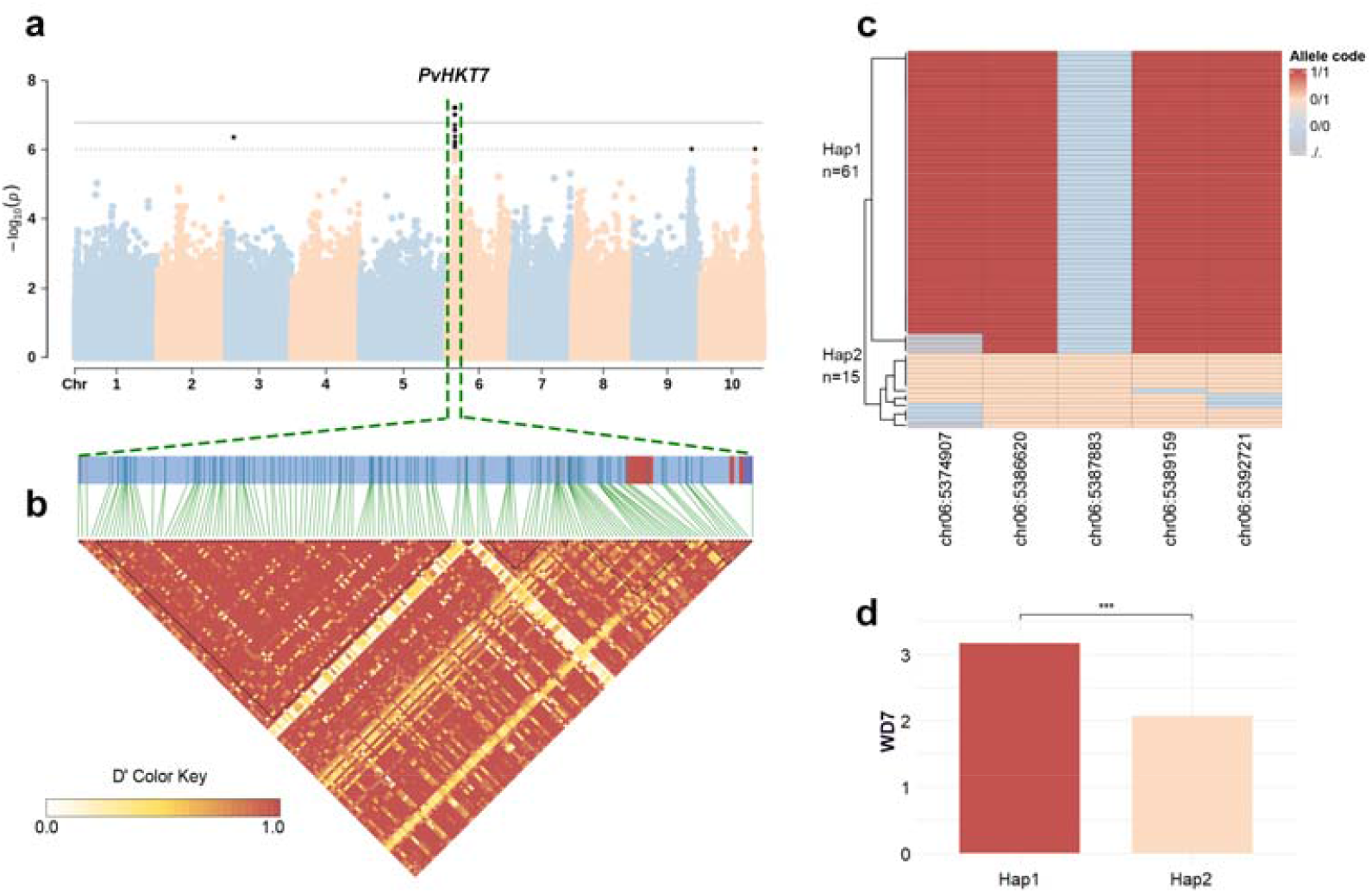
Genome-wide identification of candidate genes associated with salt tolerance. **a** Manhattan plot of the salt tolerance trait on the chromosome 6. **b** LD heat map of the *PvHKT7* gene. The color key indicates D’ values. **c** Haplotypes for the candidate gene *PvHKT7*. **d** The SNP genotype is associated with the salt tolerance phenotype.

### Selective sweep signals during salt tolerance improvement

We divided *P. vaginatum* in this study into two population, high salt tolerance population (ST) and salt sensitive population (SS) based on their phenotypes after salt stress. The nucleotide diversity (π) of the SS (1.85 × 10^−3^) was higher than ST (1.40 × 10^−3^), which indicated salt tolerance may be subject to similar selection in the course of evolution. To detect selective sweeps driven by salt environment, we compared genomic variations with high fixation index (FST) and reduction of diversity (ROD) between the ST and SS. Above the dashed horizontal thresholds of top 5%, we identified 478 overlapping sweeps between ST and SS containing 622 putative genes (Table S21). GO enrichment analysis revealed that genes in the FST and ROD overlapping were enriched in two groups of GO classifications (MF and BP), including potassium ion transmembrane transporter activity (GO:0015079), metal ion transmembrane transporter activity (GO:0046873) and xyloglucan metabolic process (GO:0010411) etc. (Fig. S8; Table S22).

### Transcriptome analysis and genetic authentication

To identify differentially expressed salt-tolerance related genes, we collected roots and leaves from accession grin_UPG145 (salt tolerance accession) at 5 time points under salt stress. The highest number of differentially expressed genes (DEGs) (Fig. S10) was detected in leaves at 12 h after salt stress, with 3379 up- and 3460 down-regulated genes. The lowest number of DEGs occurred in roots at 5D, when only 3,866 genes were differentially expressed.

Furthermore, putative *HKT* genes were found by BLASTP in *P. vaginatum*. A total of 6 candidate members in *P. vaginatum* were scanned on the basis of hidden Markov model (HMM) search (Fig. S11a). There are up-regulated, down-regulated, or even no difference in all *HKT* genes. But only *PvHKT7* was significantly up-regulated in every treatment group compared to the control group (Fig. S11b), indicating that *PvHKT7* gene plays a more important role under salt stress. Finally, we demonstrated the function of the *PvHKT7* gene in transgenic *A. thaliana* which was not harboring a *HKT7* gene itself. Three independent transgenic lines with high expression levels of *PvHKT7* (Line1, Line2, Line3) and a non-transgenic (WT) line were selected for salt tolerance evaluation. To test the effect of *PvHKT7* overexpression on salt tolerance, 3-week-old plants of Line1, Line2, and Line4 were exposed to 32 ds·m^-1^ NaCl and 1/2 MS medium for 10 days. The results showed that biomass weight of transgenic plants was significantly greater than that of WT under the salt treatment (Fig. 6). The root length of transgenic plants was longer than WT in both two type mediums. The K^+^ content in leaves and roots of transgenic plants was significantly higher than that of WT under salt treatment. These results indicated that *PvHKT7* gene enhanced salt tolerance of plants by enhancing K^+^ absorption. These results indicating overexpression of *PvHKT7* increased salt tolerance.

**Fig. 6.**
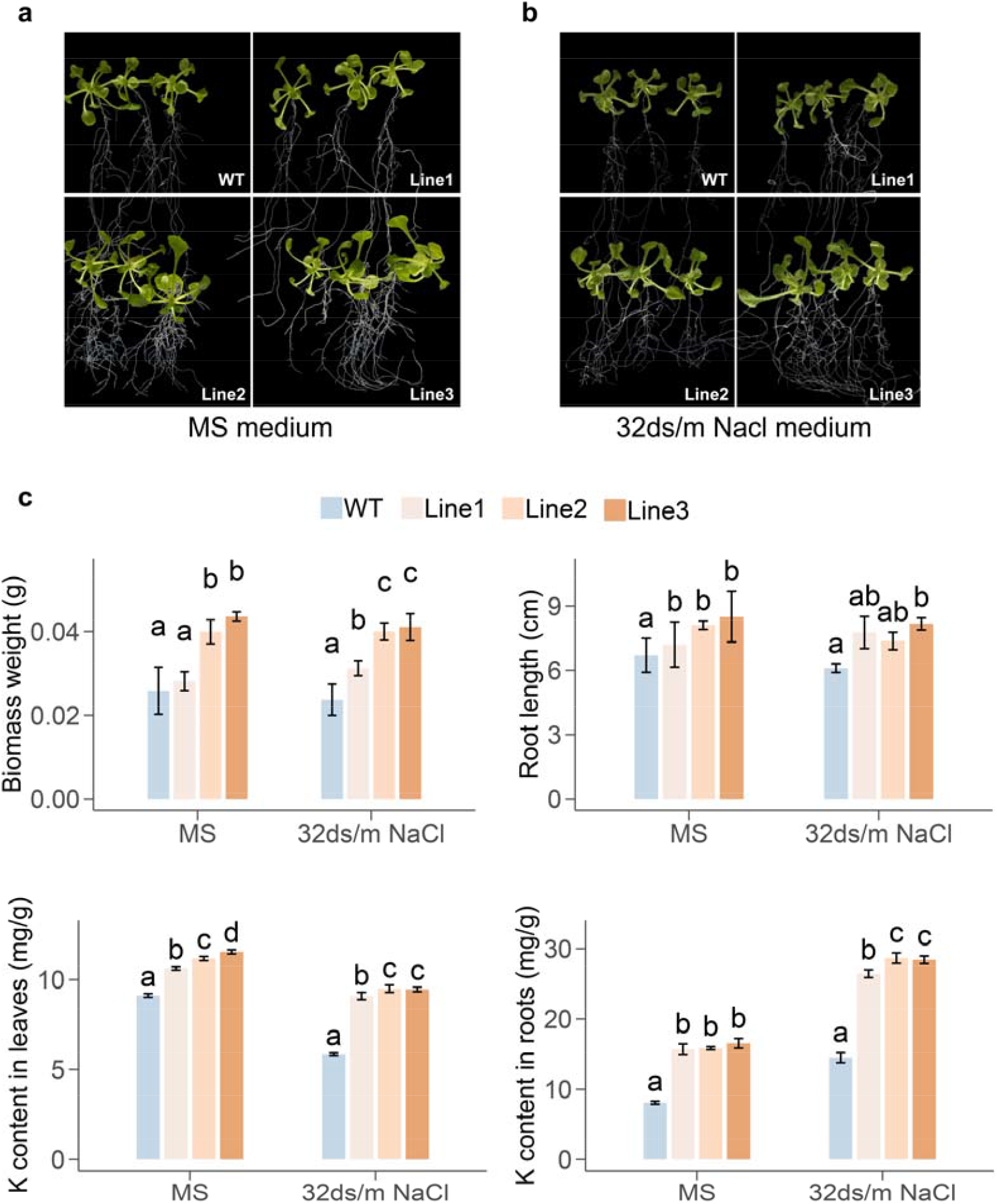
Transgenic verification. **a** The growth of non-transgenic (WT) and transgenic (Line1, Line2, Line3) *A. thaliana* exposed to 1/2 MS medium. **b** The growth of non-transgenic (WT) and transgenic (Line1, Line2, Line3) *A. thaliana* exposed to 32 ds·m^-1^ NaCl MS medium. **c** Difference analysis of biomass weight and root length between A and B non-transgenic and transgenic *A. thaliana*.

## Discussion

*P. vaginatum* is an exceptionally salt tolerant grass species that inhabits warm, coastal areas worldwide. Its ability to thrive in saline environments makes it a vital resource that could contribute as a model to the study of salt tolerance in the grasses, and as a potential source of salt tolerance genes that could be used to improve the salt tolerance of other species. However, genetic studies of *P. vaginatum* are limited by relatively few genomic resources. A highly continuous and complete reference genome is essential for a wide range of population genetics studies and experimental research. We selected the worldwide major cultivar SeaIsle2000 for *de novo* sequencing and assembled the reference genome. By combining four sequencing and assembly technologies, we proposed a high-quality chromosome-scale assembly genome with the high continuity and integrity. The SeaIsle2000 reference genome size was 517.98 Mb, including 10 chromosomes and 169 scaffolds. We evaluated the consistency and syntenic sequences with the first available published genetic maps of *P. vaginatum* and showed our chromosome assembly had good collinearity when compared with previous linkage groups.

Comparative analyses showed that *P. vaginatum* has a close genetic relationship with many grain and forage producing members of the Panicoideae, such as *S. bicolor* and *S. italica*. Comparative analyses showed that each seashore paspalum chromosome was syntenic to and highly colinear with a single sorghum chromosome. The synteny analysis indicated that *P. vaginatum, S. bicolor*, and *Z. mays* may share a common ancestor, but there was various salt tolerance among them. *P. vaginatum* is a salt tolerant grass which has an impressive level of salt tolerance and experiences its greatest productivity under salt exposure of ∼15dS m^-1^, is self-incompatible so individuals are obligate out-crossers and highly heterozygous^4^. Sorghum is believed to tolerate soil and water salinity up to 6.8 and 4.5 dS m^-1^ of electrical conductivity, respectively. Above these thresholds, a 16% yield reduction is expected per each soil salinity unit increase(Calone et al., 2020). Maize is moderately sensitive to salt stress and soil salinity is a serious threat to its production worldwide(Farooq et al., 2015). The genetic loci in seashore paspalum that confers salt tolerance can be leveraged in these important relatives in the future.

GWAS results identified several target regions that putatively control salt tolerance. Both GWAS and transcriptomes showed that *HKT7* gene was particularly important for salt tolerance in *P. vaginatum*, and the overexpression of *HKT7* increased the salt tolerance of *A. thaliana. HKT7* was not found in *A. thaliana*, which indicated that the transgenic *A. thaliana* of *HKT7* could enhanced the salt tolerance. *HKT7* is an example of the value that target gene identification can bring to future functional studies and molecular breeding. Our overarching goal in undertaking this research was to expand our understanding of seashore paspalum and to identify salt tolerance candidate genes that could be put forward for further characterization. Moreover, these data and information are valuable resources for *P. vaginatum* research and breeding, and for comparative genomic analysis of Poaceae species.

## Methods

### Sample collection

A total of 107 *P. vaginatum* accessions or cultivar genotypes with variation in morphological characteristics and geographic origin were used in this study including 24 germplasm resources collected from south China and 83 germplasm resources from United States stored in the USDA Plant Genetic Resources Conservation Unit (USDA-PCGRU) and the University of Georgia (UGA). Detailed information about the country of origin and ecotype for each accession is shown in Table S13. The world map showing origin information (Fig. 4a) was made using the R package ggplot2(Wickham, 2016). All materials are preserved as germplasm resources of *P. vaginatum* at the experimental field of Danzhou campus of Hainan University (N: 19°30′15.63″; E: 109°29′12.70″), which were respectively planted in 1 m×1 m plots.

### Genome sequencing and assembly

Genomic DNA extracted from young leaves of ‘SeaIsle2000’ were sent to Metware Metabolic Technology (Wuhan, China) for library construction and sequencing by Illumina HiSeq2000 platform, PacBio platform and 10x genomics platform, respectively. An inhouse quality control process was applied to the reads that passed the Illumina quality filters. Firstly, adapters and primer sequences were removed from the reads. Next, we discarded low-quality nucleotides (Q<30) from both ends of the reads. Then, reads with N was more than 10% were removed. Finally, When the number of low-quality (less than 5) bases in a single-end read exceeded 20%, the pair end reads were discarded too. For Pacbio long reads datas, pre-assemble reads after self-correction were used for genome assembly by Overlap-Layout-Consensus algorithm in FALCON. Then, we used Illumina reads to perform sequence error correction of contigs in Pilon(Walker et al., 2014). For 10× genomics data, the gel beads are connected with: Illumine P5 connector, 16-base Barcodes, Illumina read 1 sequencing primer and 10 bp random sequence primers. The consensus sequences were further scaffolded by integrating with 10× genomics linked reads by fragScaff software(Adey et al., 2014). Others pipelines were according to standard processes of Matware company’s standard process. Finally, all linked-reads were assist to Pacbio datas for genome assembly.

### Hi-C sequencing and analysis

The harvestable young leaves of SeaIsle2000 were fixed with formaldehyde for Hi-C sequencing. Approximately 3 g of leaf sample was collected and used for a Hi-C pipeline as described in the publication(Xie et al., 2015). The Hi-C experiments were performed by Matware company. The Hi-C libraries were then sequenced on an Illumina HiSeq PE150 platform. Hi-C reads were aligned to the raw reference genome SeaIsle2000 for high assembly quality using BWA(Li and Durbin, 2009) and LACHESIS with the default parameter settings(Burton et al., 2013).

### Assessment of genome assembly and annotation

#### Assessment of genome quality

The 1614 conserved protein models in the BUSCO niport yte_odb10 dataset and the 242 conserved protein models in the CEGMA dataset were searched against the SeaIsle2000 genome by using the BUSCO (v5.2.2)(Manni et al., 2021) and the CEGMA (v. 2.5)(Parra et al., 2007) programs with default parameters.

### Transposable element (TE) annotation

Chosen repeat libraries (RepBase; http://www.girinst.org/repbase/; Repeatmasker and repeatproteinmask; http://www.repeatmasker.org/) and *de novo* prediction were applied to identify repetitive elements. TE libraries were constructed with RepeatModeler (http://www.repeatmasker.org/) and were applied to mask the SeaIsle2000 genomes by using RepeatMasker software with default parameters (http://www.repeatmasker.org/).

### ncRNA_annotation

tRNAscan-SE (http://lowelab.ucsc.edu/tRNAscan-SE/) was used to identified tRNA in the SeaIsle2000 genome sequence. Since rRNA is highly conserved, rRNA information of SeaIsle2000 could be gained by blasting rRNA sequences of closely related species to the reference genome. The miRNA and snRNA were predicted by INFERNAL (http://infernal.janelia.org/).

### Gene prediction and function annotation

We performed an integrated approach combining PASA(Haas et al., 2003) and EVM(Haas et al., 2008) pipelines. The predicted gene models from EVM were then updated by PASA assembly alignments. Gene functions were assigned according to the best alignment using BLASTP(Altschul et al., 1997) (Evalue <10-4) to the SwissProt database (http://www.uniprot.org/), Nr database (http://www.ncbi.nlm.nih.gov/protein),Pfam database (http://pfam.xfam.org/),and the KEGG database (http://www.genome.jp/kegg/). The GO term for each gene was achieved from the corresponding InterProScan.

### Comparative Genomic and Evolutionary Analysis

#### Gene family Cluster and phylogenomic tree analysis

To identify gene family, we analyzed protein-coding genes from 10 species, *Sorghum bicolor* (*S. bicolor*), *Oryza sativa* (*O. sativa*), *Zea mays* (*Z. mays*), *Hordeum vulgare* (*H. vulgare*), *Aegilops tauschii* (*A. tauschii*), *Setaria italica* (*S. italica*), *Brachypodium distachyon* (*B. distachyon*), *Medicago truncatula* (*M. truncatula*), *Arabidopsis thaliana* (*A. thaliana*), *Paspalum vaginatum* (*P. vaginatum*). Orthologous gene groups of *P. vaginatum* and 9 other species were identified by the OrthoMCL program(Li et al., 2003). Firstly, those genes with coding protein shorter than 50 amino acids were filtered, and the longest transcript of one gene was kept when it contained more than one transcript. Then, similarity between protein sequences of all species were gained by blastp (e-value was less than 1e-5, both query and hit were more than 70% length coverage).

To infer the phylogenetic placements of *P. vaginatum*, single-copy genes of 10 species were extracted to align using MUSCLE (Edgar, 2004). The aligned results were used to infer the maximum likelihood trees with RaxML(Stamatakis, 2006). We used the mcmctree program of PAML (Yang, 2007) to estimate the divergence time among 10 species with main parameters (burn-in = 10,000, sample-number = 100,000, and sample-frequency = 2). The calibration points were selected from TimeTree website (http://www.timetree.org) as normal priors to restrain the age of the nodes,

Based on the phylogenetic tree and calibration points selected from TimeTree website, we used the ‘mcmctree’ module of PAML (http://abacus.gene.ucl.ac.uk/software/paml.html) to estimate the divergence time of these species.

### Analysis of genome synteny

The longest protein sequence of all genes was used to perform synteny searches to identify syntenic genes of *S. paspalum* versus *S. bicolor and S. paspalum* versus *Z. mays* by using BLAST(Altschul et al., 1997) and MCScanX(Wang et al., 2012). Blastp was used to search for potential anchors (E-value < 1e-10; top ten matches) between each possible pair of chromosomes in multiple genomes. Then we used the ‘circle_plotter’ and ‘bar_plotter’ of the downstream analyses programs to show the syntenic genes.

### Population Analysis

#### Population Genome Resequencing and Detection of Nucleotide Variants

All clean reads from each accession were mapped to the SeaIsle2000 reference genome using BWA (Burrows-Wheeler Aligner)(Li and Durbin, 2009) (version 0.7.17) using the mem function. Mapped reads were converted into BAM files using SAMtools(Li et al., 2009), after removing duplicate reads, SNPs and InDels within the 193 accessions variants were called by Genome Analysis Toolkit (GATK) (version 3.4-46)(McKenna et al., 2010), the raw variants were filtered using the GATK VariantFiltration tool with parameters as QD < 2.0, MQ < 30.0, FS > 60.0, SOR > 3.0, MQRankSum < -12.5, ReadPosRankSum < -8.0. Then to exclude SNP calling errors caused by incorrect mapping or InDels, a total of 3,166,505 high-quality SNPs with parameters as –max-missing 0.8 –maf 0.05 –mac 3 –minQ 30 –minDP 3 –min-alleles 2 –max-alleles 2, were kept for subsequent analysis. The identified InDels were filtered using GATK filters (QD < 2.0 || FS > 200.0 || SOR > 10.0 || MQRankSum < -12.5 || ReadPosRankSum < -8.0). SNPs/ InDels annotation was performed on the basis of the SeaIsle2000 genome with ANNOVAR (Wang et al., 2010), SNPs/ InDels were grouped into exon, intron, 5′-untranslated region and 3′-untranslated region, upstream and downstream regions (within 1 kb region from the transcription start or stop site), and intergenic regions. The SNPs/InDels in coding exons were further grouped into synonymous or nonsynonymous mutations. The SNPs causing gain of a stop codon, loss of a stop codon, or splicing were designated as large-effect SNPs. We further classified InDels in coding exons as frameshift deletions or non-frameshift deletions and the distribution of SNPs/InDels in the genome was demonstrated by Circos(Krzywinski et al., 2009).

### Population structure and phylogenetic analyses

The population genetic structure was examined using the program sNMF (v1.2) with K values from 2 to 10. We constructed a neighbor-joining tree with 3.08 million SNPs using PHYLIP software and then visualized it with the online tool iTOL (https://itol.embl.de). PCAs were done by GCTA(Yang et al., 2011). Nucleotide diversity (π) and fixation index (FST) were calculated by Vcftools(Danecek et al., 2011) with a 200kb sliding window. To estimate and compare the pattern of LD among different groups, the squared correlation coefficient (r^2^) between pairwise SNPs was computed and plotted using the PopLDdecay (v.3.40)(Zhang et al., 2019) software. Parameters in the program were MaxDist 500. The average r^2^ value was calculated for pairwise markers in a 500-kb window and averaged across the whole genome.

### Phenotypic data for salt tolerance

After treatment with salt solution, the symptoms of salt damage in the leaves were assessed visually. Seventy-six germplasms of *P. vaginatum* were chosen from the lawn grass germplasm garden, all materials were propagated from healthy stolons gathered from their native habitat and planted in the experimental field of Danzhou campus of Hainan University. From July 15 to November 25 in 2019, each variety was acclimatized from the same number of tillers, planted in four 5.5×5.5 cm circular containers and place on gauze-based foam board (30 cm×30 cm), with hydroponic culture conditions using Hoagland’s nutrient medium in a greenhouse with an average temperature from 28 °C to 36 °C. The Hoagland’s nutrient solution was replaced once every week for about ten weeks until each circular container was covered with grass and had the same growth state. The experimental design was a randomized block design with 2 treatments (control and salt treatment), and 3 replications for each treatment. At first, salt treatment was performed with 32 ds·m^-1^ NaCl for 3 days, the visual reactions of plant under salt stress were evaluated using wilting degree (WD3) and withering rates (WR3). Then NaCl was added to 54 ds·m^-1^ until the 7th day, wilting degree (WD7) and withering rates (WR7) were measured again.

### FST and ROD analysis and selective sweep detection

The top 20 salt-tolerant germplasms were named ST while the most 20 sensitive-salt germplasms were named ST according to their phenotype after salt stress. Nucleotide diversity (π), fixation index (FST) and reduction of diversity ROD were calculated by Vcftools(Danecek et al., 2011) with a 200 kb sliding window with a step size of 20 kb. ggplot2(Wickham, 2016) in the R packages was applied for presentation.

### Genome-wide association study

Only SNPs with MAF≥0.05 and missing rate ≤ 0.2 in a population were used to carry out GWAS. This resulted in 2,221,123 SNPs that were used in GWAS for 76 *P. vaginatum* germplasms. We performed GWAS using EMMAX software. Significant *P*-value (0.05/n, Bonferroni correction) and suggestive *P*-value (10^−6^) were set to control the genome-wide type I error rate. The results of GEMMA were visualized as Manhattan and Q-Q plots with the R package ‘Cmplot’. We then selected genes near the peak SNPs according to the Manhattan plot for each trait and annotated candidate genes.

### GO terms and KEGG pathway enrichment analysis

Based on the annotation information (http://eggnog-mapper.embl.de/)(Huerta-Cepas et al., 2017), candidate genes were analyzed by GO terms and KEGG pathway functional enrichment using clusterProfiler under the R platform (v4.0.2). GO terms and KEGG pathways with Q-values < 0.05 were considered to be significantly enriched.

### Transcriptome analysis and genetic authentication

Total RNA was isolated from a sampled organ with three biological replicates at different stress stages to investigate expression of the genes associated with salt tolerance for roots and leaves. RNA extraction and library preparation for each sample were performed by Matware company. All 64 samples with three biological replicates were sequenced using the Illumina HiSeq 2000 platform, and 150 bp pairedend reads were generated. After filtering, 2,464,855,778 clean reads were obtained, containing 369.73 Gb of data. On average, 88.37% of the reads uniquely mapped to the SeaIsle2000 reference genome (Table S23). Analysis of differential gene expression between two samples was performed using the DESeq R package (v1.18.0). Genes with an adjusted P value <0.05 found by DESeq were assigned as differentially expressed.

*A. thaliana* was used for transformation in the present study. To generate *PvHKT7* overexpression lines, a 1806-bp coding sequence (CDS) of *PvHKT7* was amplified from cDNA of SeaIsle2000 and verified by sequencing, using the primer set listed in Supplementary Table S27. The gene entry vector and expression vector were constructed by enzyme digestion ligand and Plasmid DNA was extracted using a plasmid microextraction kit (MEgi bio). Transgenic *A. thaliana* seeds of T2 generation and WT generation were cultured on LB medium plate (32 ds·m^-1^ NaCl and normal) at 25 °C for 10 days

## Contributions

Li Liao, Xu Hu, Jiangshan Hao and Minqiang Tang contributed equally to this work. Jie Luo and Zhiyong Wang conceived and managed the project. Li Liao, Shangqian Xie and Minqiang Tang designed the experiments. Longzhou Ren performed molecular cloning and salt stress experiments. Xu Hu and Jiangshan Hao performed data analyses and wrote the manuscript. Ling Pan, Paul Raymer, Peng Qi and Zhenbang Chen interpreted the results and revised the manuscript.

## Data available

The raw sequence data of *P. vaginatum* genome project has been deposited at the The National Center for Biotechnology Information under BioProject PRJNA848273. The final assembly and gene annotation of *P. vaginatum* is available at GenBank under the accession number SUB11601520.

## Acknowledgements

We sincerely acknowledge the Katrien M. Devos, which is a professor of University of Georgia, for a critical reading of the manuscript and valuable discussions. This work was supported by the National Natural Science Foundation of China (No.32060409), the Construction of World First Class Discipline of Hainan University (No.RZZX201905) and National Project on Sci-Tec Foundation Resources Survey (2017FY100600).

## Supplementary Information

Supplementary Fig.

Supplementary Table

